## Supplementary Table 11 for "*FSH*β links photoperiodic signalling to seasonal reproduction in Japanese quail"

### Demultiplexing, basecalling
for dir in RawSeqData/*/
do
dir1=${dir%*/}       # remove the trailing "/"
base1=${dir1##*/}    # print everything after the final "/" >>>> This is the run number
guppy_basecaller -i RawSeqData/${base1}/fast5/ -s RawSeqData/${base1}/ --flowcell FLO-MIN106 --kit SQK-DCS109 --barcode_kits EXP-NBD104 --trim_barcodes -x "cuda:0"
	for bardir in RawSeqData/${base1}/barcode*/
	do
	dir2=${bardir%*/}      # remove the trailing "/"
	base2=${dir2##*/}      # print everything after the final "/" >>>> This is the barcode number
	cat RawSeqData/${base1}/${base2}/*.fastq > RawData/${base1}_${base2}.fastq
	done
done


### trimming, and filtering 
mkdir RawData/temp/
for infile in RawData/*.fastq
do
base=$(basename ${infile} .fastq)
porechop -i RawData/${base}.fastq -o RawData/temp/${base}.fastq --extra_end_trim 20 -t 22
filtlong --min_length 25 --target_bases 5000000000000 --mean_q_weight 9 RawData/temp/${base}.fastq > ./RawData/${base}.fastq
done
rm -r ./RawData/temp/


### Main loop ###
### index ref
minimap2 -t 22 -I 1000G -d ./ReferenceData/transcript.mmi ./ReferenceData/transcript.fna

mkdir Analysis/
mkdir Analysis/Minimap/
mkdir Analysis/samtools/
mkdir Analysis/Salmon/
mkdir Analysis/Results/

for infile in ./RawData/*.fastq
do
base=$(basename ${infile} .fastq)
# align to ref
minimap2 -t 22 -ax map-ont -p 1.0 -N 100 ./ReferenceData/transcript.mmi ${infile} | samtools view -Sb > ./Analysis/Minimap/raw${base}.bam
# sort alignments
samtools sort ./Analysis/Minimap/raw${base}.bam -o ./Analysis/Minimap/${base}.bam -@ 22
rm ./Analysis/Minimap/raw${base}.bam
# index alignments
samtools index ./Analysis/Minimap/${base}.bam
# some useful stats
samtools flagstat ./Analysis/Minimap/${base}.bam > ./Analysis/samtools/${base}.flagstat
# count reads
salmon quant --noErrorModel -p 22 -t ./ReferenceData/transcript.fna -g ./ReferenceData/annotation.gff -l SF -a ./Analysis/Minimap/${base}.bam -o ./Analysis/Salmon/${base}
done

# QC report scipt
R --slave -e 'rmarkdown::render("NanoFlow_QC_Report.Rmd", "html_document")'

# analysis scripts....the report ones stopped working when called from the commandline for some reason!!!
# Still work if you load them in R studio, set WD, then knit.
#R --slave -e 'rmarkdown::render("NanoFlow_Report_NoIso.Rmd", "html_document")'
#R --slave -e 'rmarkdown::render("NanoFlow_Report.Rmd", "html_document")'


NanoFlowreport.rmd
---
title: "NanoFlow: cDNA Transcriptome Report"
date: 'Report created: `r Sys.Date()`'
#bibliography: Static/Bibliography.bib
output:
  html_document:
    css: Static/UoG.css
    df_print: paged
    highlight: null
    keep_md: yes
    number_sections: yes
    self_contained: yes
    theme: default
    toc: yes
    toc_depth: 2
    toc_float:
      collapsed: yes
      smooth_scroll: yes
  word_document:
    toc: yes
    toc_depth: '2'
link-citations: yes
always_allow_html: yes
---

<div style="position:absolute;top:0px;right:0px;width:35%;">
```{r, echo=FALSE}
knitr::include_graphics("Static/Images/UoG.png")
```
</div>


```{r Imports, echo=FALSE, results='hide', warning=FALSE, message=FALSE}
library(grid)
library(gridExtra)
library(pcaMethods)
library(yaml)
library(DRIMSeq)
library(AnnotationDbi)
library(GenomicFeatures)
library(dplyr)
library(edgeR)
library(DEXSeq)
library(DESeq2)
library(devtools)
library(ggplot2)
library(tidyr)
library(reshape2)
#library(session)     
library(writexl)
library(digest)
library(kableExtra)
library(stageR)
library(ShortRead)  # Oh, the irony!

config <- yaml.load_file("config.yaml")

sig_level <- config$adjPValueThreshold
gffFile <- file.path("ReferenceData", config$annotation)   
# Format set as gtf as auto wasn't working. See line 152 to change.

resultDir <- file.path("Analysis", "Results")
dir.create(resultDir, showWarnings = FALSE, recursive=TRUE)
persistenceData <- file.path(resultDir, "NanoFlow_Report.Rdata")

```

# Study design

The **`NanoFlow_Report_NoIso.Rmd`** script has been used to generate this report. The supplied **`config.yaml`** configuration file defines the biological samples used in this DGE and DTU analysis.

```{r setupParameters, echo=FALSE}
studyDesign <- data.frame()
for (i in 1:length(config$Samples)) {
  studyDesign <- rbind(studyDesign, 
                       data.frame(samples=names(config$Samples[[i]][[1]]), 
                                  filename=unlist(config$Samples[[i]][[1]]), 
                                  group=names(config$Samples[[i]])))
}

studyDesign$replicate <- sapply(1:nrow(studyDesign), function(x)sum(studyDesign$group[1:x]==studyDesign$group[x]))
# reorder for stated reference group
studyDesign$group <- relevel(as.factor(studyDesign$group), ref=config$referenceGroup)
# quick tidy
studyDesign <- studyDesign[,-which(colnames(studyDesign)=="samples")]

knitr::kable(studyDesign, booktabs=TRUE, table.envir='table*', linesep="")  %>%
  kable_styling(latex_options=c("hold_position", "scale_down"))

```

*Table showing the cDNA sequence files evaluated for DGE and DTU analysis.*

```{r, echo=FALSE}
# retain file name prefix
samples <- gsub("\\..+", "", basename(as.character(studyDesign$filename)))
# this code block to replace the col_data export and import
coldata <- data.frame(cbind(NULL, sample_id=rownames(studyDesign), group=as.character(studyDesign$group), type="single-read"), stringsAsFactors=TRUE)


```

# Analysis file output

The **`Salmon`** tool (@Salmon2017) has been used to assign cDNA read counts to the annotated transcripts.

```{r importSalmonCounts, results='hide', echo=FALSE}
salmonCountFiles <- paste(file.path("Analysis","Salmon",samples,"quant.sf"))

rawSalmonCounts <- data.frame(Reference=character(), stringsAsFactors = FALSE)
harvestSalmonCounts <- function(file) {
  obs <- read.table(file, header=TRUE, stringsAsFactors = FALSE)
  rawSalmonCounts <<- full_join(rawSalmonCounts, obs[which(obs$NumReads>0), c("Name", "NumReads")], by=c("Reference"="Name"))
  file
}
lapply(salmonCountFiles, harvestSalmonCounts)
colnames(rawSalmonCounts) <- append("Reference", samples)

rownames(rawSalmonCounts) <- gsub("\\.+", "", as.character(rawSalmonCounts$Reference))            
#taken out second . in gsub conditions
```

```{r geneCounts, echo=FALSE, warning=FALSE}
# adjust column names for the presented table and excel file
vRawSalmonCounts <- rawSalmonCounts[order(rowSums(rawSalmonCounts[, seq(2, ncol(rawSalmonCounts))]), decreasing=TRUE),]
newnames <- rownames(studyDesign)[match(colnames(vRawSalmonCounts)[which(colnames(vRawSalmonCounts) %in% samples)], samples)]
colnames(vRawSalmonCounts)[which(colnames(vRawSalmonCounts) %in% samples)] <- newnames

csvExpressedTranscripts <- file.path(resultDir, "ExpressedTranscripts.csv")
write.csv(vRawSalmonCounts, file = csvExpressedTranscripts)
```

The complete transcript mapping data is provided as a CSV format file at

~~~
`r csvExpressedTranscripts`
~~~

```{r loadAnnotations, echo=FALSE, results='hide', warning=FALSE, message=FALSE}
# load the database annotation from provided GFF file
txdb <- makeTxDbFromGFF(file=gffFile, format=c("gff"))
txdf <- AnnotationDbi::select(txdb, keys(txdb,"GENEID"), "TXNAME", "GENEID")
tab <- table(txdf$GENEID)
txdf$ntx<- tab[match(txdf$GENEID, names(tab))]

### ONLY FOR THIS MOUSE REFERENCE ###
vRawSalmonCounts$Reference <- sub(".*?-","",vRawSalmonCounts$Reference)
rownames(vRawSalmonCounts) <- sub(".*?-","",rownames(vRawSalmonCounts))

# FIXME: filter for transcripts which are in the annotation. Why they are not all there? 
filteredSalmonCounts <- vRawSalmonCounts[which(vRawSalmonCounts$Reference %in% txdf$TXNAME),]

# Create counts data frame:
salmonCounts<-data.frame(txdf[match(filteredSalmonCounts$Reference, txdf$TXNAME), c(1,2)], filteredSalmonCounts)

salmonCounts[is.na(salmonCounts)] <- 0
colnames(salmonCounts) <- append(c("gene_id", "feature_id"),colnames(filteredSalmonCounts))

# adjust column names for the presented table and excel file
vSalmonCounts <- salmonCounts[order(rowSums(salmonCounts[, seq(4, ncol(salmonCounts))]), decreasing=TRUE),]
newnames <- rownames(studyDesign)[match(colnames(vSalmonCounts)[which(colnames(vSalmonCounts) %in% samples)], samples)]
colnames(vSalmonCounts)[which(colnames(vSalmonCounts) %in% samples)] <- newnames

csvExpressedGenesTranscripts <- file.path("Analysis","Results","ExpressedGenesTranscripts.csv")
write.csv(vSalmonCounts, file = "Analysis/Results/ExpressedGenesTranscripts.csv")

```

The complete transcript mapping counts and parental gene assignments is provided as a CSV format file at 

~~~
`r csvExpressedGenesTranscripts`
~~~

**`r round(as.numeric((table(table(salmonCounts$gene_id)) / length(table(salmonCounts$gene_id)) * 100)[1]), digits=1)` %** of genes contained a single expressed isoform, and the largest number of transcripts associated with a single gene is **`r max(table(salmonCounts$gene_id))` ** transcripts.

<!-- {r geneTranscriptCount, echo=FALSE}
plotData(dmDSdata(counts=salmonCounts, samples = coldata))

*Frequency plot showing the number of transcripts assigned to genes across the sampled cDNA sequence collection*
-->

Transcript count data was filtered using **`DRIMSeq`** (@R-DRIMSeq), according to filtering parameters defined in **`config.yaml`**.

```{r, echo=FALSE, warning=FALSE}
dmDSdataObj  <- dmDSdata(counts=salmonCounts, samples=coldata)
trs_cts_unfiltered <- counts(dmDSdataObj)

# Sum transcript counts into gene counts:
trs_cts <- counts(dmDSdataObj)
names(trs_cts)[2]<-"transcript_id"  # changed from featureID

gene_cts <- trs_cts_unfiltered %>% dplyr::select(c(1, 3:ncol(trs_cts)))  %>% group_by(gene_id) %>% summarise_all(list(sum)) %>% data.frame()

xlsExpressedGeneCounts <- file.path(resultDir, "FilteredExpressedGeneCounts.xlsx")
write_xlsx(gene_cts, path = xlsExpressedGeneCounts)

rownames(gene_cts) <- gene_cts$gene_id
gene_cts$gene_id <- NULL
keep = filterByExpr(gene_cts, design = design, min.prop = 0.5, min.count = 2, min.total.count = 15)
gene_cts <- gene_cts[keep,]
# write out these feature level (isoform) and gene count data to Excel files


xlsExpressedFeatureCounts <- file.path(resultDir, "FilteredExpressedFeatureCounts.xlsx")
write_xlsx(trs_cts, path = xlsExpressedFeatureCounts)

```

Filtered gene counts and gene isoform counts have been written to the files indicated below

~~~
`r xlsExpressedGeneCounts`
`r xlsExpressedFeatureCounts`
~~~

# Differential gene expression analysis

Statistical analysis performed using **`edgeR`** (@R-edgeR2010, @R-edgeR2012) on filtered gene counts, using the `TMM` method for normalisation, and correcting for false discovery rate (*FDR*) using the method of Benjamini & Hochberg (@BH1995).


```{r, echo=FALSE, results='hide', warning=FALSE, error=FALSE}
# Building model matrix - ensuring that the config.yaml specified reference is key in specified factor ...


Group <- factor(studyDesign$group, 
                levels=append(config$referenceGroup,
                              levels(studyDesign$group)[which(levels(studyDesign$group)!=config$referenceGroup)]))

design <- model.matrix(~Group)
# Differential gene expression using edgeR:

y <- DGEList(gene_cts)
y <- calcNormFactors(y)
y <- estimateDisp(y,design)
fit <- glmQLFit(y,design)
norm_counts <- cpm(y) # to plot against

numberofgroups <- length(unique(studyDesign$group))
if(numberofgroups == 2){(qlf <<- glmQLFTest(fit))
}else{
    (qlf <<- glmQLFTest(fit, coef =2:numberofgroups))}

edger_res <- topTags(qlf, n=nrow(y), sort.by="PValue")[[1]]

xlsEdgeRDiffExpr <- file.path(resultDir, "EdgeRDiffExpression.xlsx")
write_xlsx(as.data.frame(edger_res), path = xlsEdgeRDiffExpr)

myCPM <- cpm(gene_cts)

xlsNormCounts <- file.path(resultDir, "NormCPM.xlsx")
write_xlsx(as.data.frame(myCPM), path = xlsNormCounts)
```

The results of the differential expression analysis have been written to:

~~~
`r xlsEdgeRDiffExpr`
~~~

```{r diffExprGeneTable, echo=FALSE}
knitr::kable(edger_res[1:15,], digits = c(2,2,2,45,45), booktabs=TRUE, table.envir='table*', linesep="")  %>%
  kable_styling(latex_options=c("hold_position"), font_size=9)
```
*Table showing the top 15 genes, ranked by adjusted p-value, from the edgeR analysis. 
**`logFC`** = log2 fold change between experimental conditions.
**`logCPM`** = log2-scaled counts per million measure of abundance.*

```{r quick calcs, echo=FALSE}
NumUpReg <- edger_res %>% 
  filter(edger_res[,1:(numberofgroups-1)]>config$lfcThreshold,
         PValue<config$adjPValueThreshold)
NEGlfcThreshold <- config$lfcThreshold
NEGlfcThreshold[sapply(NEGlfcThreshold, is.numeric)] <- NEGlfcThreshold[sapply(NEGlfcThreshold, is.numeric)] * -1
NumDownReg <- edger_res %>% 
  filter(edger_res[,1:(numberofgroups-1)]<NEGlfcThreshold,
         PValue<config$adjPValueThreshold)

NumUpRegFDR <- edger_res %>% 
  filter(edger_res[,1:(numberofgroups-1)]>config$lfcThreshold,
         FDR<config$fdrThreshold)
NumDownRegFDR <- edger_res %>% 
  filter(edger_res[,1:(numberofgroups-1)]<NEGlfcThreshold,
         FDR<config$fdrThreshold)
```
**`r nrow(NumUpReg) + nrow(NumDownReg)`** genes were differentially expressed, with a log2 fold change threshold of *`r config$lfcThreshold`* and P-value threshold of *`r config$adjPValueThreshold`*. **`r nrow(NumUpReg)`** genes had increased expression, and **`r nrow(NumDownReg)`** genes had reduced expression.


**`r nrow(NumUpRegFDR) + nrow(NumDownRegFDR)`** genes were differentially expressed, with a log2 fold change threshold of *`r config$lfcThreshold`* and FDR threshold of *`r config$fdrThreshold`*. **`r nrow(NumUpRegFDR)`** genes had increased expression, and **`r nrow(NumDownRegFDR)`** genes had reduced expression.

```{r geneOfInterest, echo=FALSE, comment=NA} 
# Filter up/down regulated genes
UpReg <- row.names(NumUpReg[1:20,])
UpRegGenes <- as.data.frame(norm_counts[which(row.names(norm_counts)%in%UpReg),], order = row.names(NumUpReg[1:20,]))
DownReg <- row.names(NumDownReg[1:20,])
DownRegGenes <- as.data.frame(norm_counts[which(row.names(norm_counts)%in%DownReg),], order = row.names(NumDownReg[1:20,]))

# bind and transpose and rownames
UpDownGenes <- rbind(DownRegGenes, UpRegGenes)
UpDownGenes <- t(UpDownGenes)
rownames(UpDownGenes) <- as.character(studyDesign$samples)
UpDownGenes <- as.data.frame(UpDownGenes)

# add sample and group ids to melt data
UpDownGenes$sample <- as.character(row.names(UpDownGenes))
row.names(UpDownGenes) <- NULL
UpDownGenes$group <- studyDesign$group
UDGmelt <- melt(UpDownGenes, id.vars=c("sample", "group"), variable.name="gene")

# Plot
UpDownPlot <- 
  ggplot(data = UDGmelt, aes(x=gene, y=value, fill=group)) + 
  geom_boxplot() + 
  ggtitle("Top Differentially Expressed Genes") + 
  xlab("") +ylab("Norm Read Count") +
  theme_bw() +
  theme(axis.text.x = element_text(angle = 90, size=8))

```

Shown below are results from **`edgeR`** analysis: The top 20 up and 20 down differentially expressed genes are shown below.

```{r Diff Exp Plot, echo=FALSE}

print(UpDownPlot)
```

# Princliple Component Analysis

Principal Component Analysis (PCA). Principal components are calculated that describe decreasing amounts of the total explainable variation. PCA performed by **`edgeR`**.

```{r pca, echo=FALSE, include=TRUE, fig.margin=FALSE, fig.fullwidth = FALSE, cache=FALSE}

pcaMatrix <- counts(dmDSdataObj)[,-c(1:2)]
md <- prep(t(pcaMatrix), scale="none", center=TRUE)
pca <- pca(md, method="svd", center=TRUE, nPcs=3)
xdata <- as.data.frame(pca@scores)
xdata <- cbind(xdata, group=studyDesign$group)
x <- 1
y <- 2
xpercent <- round(pca@R2[x]*100, digits=1)
ypercent <- round(pca@R2[y]*100, digits=1)
xlab <- paste("Prin.Comp. ",x," (",xpercent,"%)",sep="")
ylab <- paste("Prin.Comp. ",y," (",ypercent,"%)",sep="")
PCAplot <- ggplot(xdata, aes(x=PC1, y=PC2)) + 
  geom_point(aes(fill=group), colour="black", pch=21, size=5) + 
  scale_colour_brewer(palette="Paired") + 
  geom_vline(xintercept=0, color="darkgray") + 
  geom_hline(yintercept=0, color="darkgray") + 
  ggtitle("PCA analysis of experimental samples") + 
  ylab(paste("PC2 (",ypercent,"%)",sep=""))  + 
  xlab(paste("PC1 (",xpercent,"%)",sep="")) +
  theme_bw()
print(PCAplot)
```

*The figure \ref{fig:pca} above is a PCA plot showing the distribution of sample data for the first two principal components. The first principal component is shown on the x-axis; the second on the y. The total amount of variation explained is shown on the axis legends.*

# GO and KEGG Pathway Analysis

```{r NOT WORKING GO and KEGG, echo=FALSE, warning=FALSE}
GOlist <- edger_res[which(edger_res$FDR<=config$fdrThreshold),]
GOlist <- c(row.names(GOlist))
xlsGOlist <- file.path(resultDir, "xlsGOlist.xlsx")
write_xlsx(as.data.frame(GOlist), path = xlsGOlist)

GeneList2 <-data.frame(salmonCounts[match(GOlist, salmonCounts$GENEID), 1:2])
xlsGOlist2 <- file.path(resultDir, "xlsGOlist2.xlsx")
write_xlsx(as.data.frame(GeneList2), path = xlsGOlist2)

#GOanalysis <- goana(qlf)
#capture.output(summary(mylist), file = "My New File.txt")


#GOtoptags <- topTags(qlf, n = )
#GOtoptags
#goTOP <-  goana(qlf, species.KEGG="oas")
#keg <- kegga(qlf, species.KEGG="oas")
```

A list of differentially expressed genes for pathway analysis has been written to:

~~~
`r xlsGOlist`
~~~


# Reproducible research

This report has been created for reproducibility, using **`Rmarkdown`**, publicly available **`R`** packages, and the **`LaTeX`** document typesetting software. For clarity the **`R`** packages used, and their versions, are listed below.

\fontsize{8}{12}

```{r sessionInfo, eval=TRUE, echo=FALSE, comment=NA}
options(width = 100)
utils:::print.sessionInfo(sessionInfo()[-7], locale=FALSE)
```


\fontsize{10}{14}


```{!r, engine='bash', echo=FALSE, comment=NA}
conda list "porechop|filtlong|salmon|samtools|minimap2|openjdk|rstudio"
```

The session data produced in the production of this report, which can be used for further analysis of the dataset, can be found here:

~~~
`r persistenceData`
~~~

```{r, echo=FALSE, include=FALSE, warning="hide"}
save.session(persistenceData)
```


\pagebreak


# References and citations

Unfortunately due to package changes that's caused an issue with my code, this section is not working right now.
If you need the citations, then please open the file `Static/Bibliography.bib` with a text editor.


Heatmap

install.packages("BiocManager")
install.packages("tidyverse")
#need r version 4.1.0 for this to function
if (!requireNamespace("BiocManager", quietly = TRUE))
  install.packages("BiocManager")
BiocManager::install(version = "3.13")
BiocManager::install("ComplexHeatmap")


library(ComplexHeatmap)
library(rstudioapi)
library(tidyverse)
library(ggplot2)
library(grid) 
library(RColorBrewer) 
library(gdata) 
library(plots) 
library(reshape2)
library(circlize)
library(ggstatsplot)
library(pheatmap)
library(readxl)

setwd(dirname(getActiveDocumentContext()$path))
z_scores <- read_excel("z_scores.xlsx")

#If you have to transpose first
#z_scores = as.data.frame(t(Quial_zscores_ordered))

#config, set colours and heatmap size
my_palette <- colorRamp(c(-2,0,2), c("red", "yellow", "green"))(n = 299)
number = 399

heat = z_scores
row.names(heat) = heat$Label
names = row.names(heat)
heatdata = heat[, -1]
row.names(heatdata) = row.names(heat)


# fix order of the clusters to have 1 to 4, top to bottom
#pamClusters$clustering <- factor(pamClusters$clustering,
#levels = c('Cluster 1', 'Cluster 2', 'Cluster 3', 'Cluster 4'))

heatdata = heatdata[0:number,]
names = names[1:number]
row.names(heatdata) = names
heatdata = as.matrix(heatdata)
colnames(heatdata) = factor(colnames(heatdata), levels = c("DL16A", "DL14A", "DL12A", "DL10A", "DL8", "DL10V", "DL12V", "DL14V", "DL16V"))
pamClusters <- cluster::pam(heatdata, k = 10) # pre-select k = n centers
Clusters <- cluster::pam(heatdata, k = 10)
pamClusters1 <- fpc::pamk(heatdata)
pamClusters$clustering <- paste0('', pamClusters$clustering)


# fix order of the clusters to have 1 to 4, top to bottom
#pamClusters$clustering <- factor(pamClusters$clustering,
#levels = c('Cluster 1', 'Cluster 2', 'Cluster 3', 'Cluster 4'))


h = Heatmap(heatdata, 
            name = "Z Score",
            col=colorRamp2(c(-1, 0, 1), c("blue", "white", "red")),
            show_row_names = F,
            row_names_gp = gpar(fontsize = 10),
            column_order = c("DL16A", "DL14A", "DL12A", "DL10A", "DL8", "DL10V", "DL12V", "DL14V", "DL16V"),
            #row_ha = rowAnnotation(foo = runif(n), bar1= anno_barplot(n)),
            #right_annotation = boxplotRow,
            row_split = factor(pamClusters$clustering, levels = c(1, 10, 9, 2, 6, 5, 7, 8, 3, 4)),
            cluster_row_slices = F,
            #clustering_distance_rows = "kendall",
            row_dend_reorder = F
)


draw(h)


---
title: "WGCNA"
output:
  pdf_document: default
  html_document: default
---


```{r setup, include=FALSE}
knitr::opts_chunk$set(fig.path = "figures/")
```

## R Markdown

This is an R Markdown document. Markdown is a simple formatting syntax for authoring HTML, PDF, and MS Word documents. For more details on using R Markdown see <http://rmarkdown.rstudio.com>.

When you click the **Knit** button a document will be generated that includes both content as well as the output of any embedded R code chunks within the document. You can embed an R code chunk like this:

## WGCNA
This script has been written to perform WGCNA analyses on hamster and quail seasonal sequencing data.

## Input and Cleaning
You only need to run this chunk once for each analysis, future chunks will automatically load necessary data.

## IMPORTANT!!!!
check the cut-off for exclusion is appropriate for your data. This script will not work if you do not!
You 


```{r input, echo=FALSE}
library(WGCNA)
library(rstudioapi)
library(tinytex)
library(edgeR)

setwd(dirname(getActiveDocumentContext()$path))


# The following setting is important, do not omit.
options(stringsAsFactors = FALSE);
#Read in the female liver data set
femData = read.csv("CPM.csv");
# Take a quick look at what is in the data set:
dim(femData);
names(femData);


datExpr0 = as.data.frame(t(femData[, -1]));
names(datExpr0) = femData$Gene;
#rownames(datExpr0) = names(femData)[, -1];

#datExpr0 = as.data.frame(t(scale(as.data.frame(t(datExpr0)))))

gsg = goodSamplesGenes(datExpr0, verbose = 3);
gsg$allOK


if (!gsg$allOK)
{
  # Optionally, print the gene and sample names that were removed:
  if (sum(!gsg$goodGenes)>0) 
    printFlush(paste("Removing genes:", paste(names(datExpr0)[!gsg$goodGenes], collapse = ", ")));
  if (sum(!gsg$goodSamples)>0) 
    printFlush(paste("Removing samples:", paste(rownames(datExpr0)[!gsg$goodSamples], collapse = ", ")));
  # Remove the offending genes and samples from the data:
  datExpr0 = datExpr0[gsg$goodSamples, gsg$goodGenes]
}


sampleTree = hclust(dist(datExpr0), method = "average");
# Plot the sample tree: Open a graphic output window of size 12 by 9 inches
# The user should change the dimensions if the window is too large or too small.
sizeGrWindow(12,9)
#pdf(file = "Plots/sampleClustering.pdf", width = 12, height = 9);
par(cex = 0.6);
par(mar = c(0,4,2,0))
plot(sampleTree, main = "Sample clustering to detect outliers", sub="", xlab="", cex.lab = 1.5, 
     cex.axis = 1.5, cex.main = 2)


# Plot a line to show the cut
abline(h = 25000, col = "red");
# Determine cluster under the line
clust = cutreeStatic(sampleTree, cutHeight = 25000, minSize = 10)
table(clust)
# clust 1 contains the samples we want to keep.
keepSamples = (clust==1)
datExpr = datExpr0[keepSamples, ]
nGenes = ncol(datExpr)
nSamples = nrow(datExpr)

plot(sampleTree, main = "Sample clustering to detect outliers", sub="", xlab="", cex.lab = 1.5, cex.axis = 1.5, cex.main = 2)
abline(h = 25000, col = "red");
# Determine cluster under the line
clust = cutreeStatic(sampleTree, cutHeight = 25000, minSize = 10)
table(clust)
# clust 1 contains the samples we want to keep.
keepSamples = (clust==1)
datExpr = datExpr0[keepSamples, ]
nGenes = ncol(datExpr)
nSamples = nrow(datExpr)


traitData = read.csv("Physio.csv", fileEncoding="UTF-8-BOM");
dim(traitData)
names(traitData)
# remove columns that hold information we do not need.
allTraits = traitData[];
allTraits = allTraits[];
dim(allTraits)
names(allTraits)
# Form a data frame analogous to expression data that will hold the clinical traits.
femaleSamples = rownames(datExpr);
traitRows = match(femaleSamples, allTraits$Mice);
datTraits = allTraits[traitRows, -1];
rownames(datTraits) = allTraits[traitRows, 1];
collectGarbage();


# Re-cluster samples
sampleTree2 = hclust(dist(datExpr), method = "average")
# Convert traits to a color representation: white means low, red means high, grey means missing entry
traitColors = numbers2colors(datTraits, signed = FALSE);
# Plot the sample dendrogram and the colors underneath.


plotDendroAndColors(sampleTree2, traitColors,
                    groupLabels = names(datTraits),
                    main = "Sample dendrogram and trait heatmap")


save(datExpr, datTraits, file = "input.RData")

```


```{r network construction, echo=FALSE}

# Display the current working directory
getwd();
# If necessary, change the path below to the directory where the data files are stored.
# "." means current directory. On Windows use a forward slash / instead of the usual \.
workingDir = ".";
setwd(workingDir);
# Load the WGCNA package
library(WGCNA)
# The following setting is important, do not omit.
options(stringsAsFactors = FALSE);
# Allow multi-threading within WGCNA. This helps speed up certain calculations.
# At present this call is necessary for the code to work.
# Any error here may be ignored but you may want to update WGCNA if you see one.
# Caution: skip this line if you run RStudio or other third-party R environments.
# See note above.
enableWGCNAThreads()
# Load the data saved in the first part
lnames = load(file = "input.RData");
#The variable lnames contains the names of loaded variables.
lnames


```

```{r scale-free topology, echo=FALSE}

# Choose a set of soft-thresholding powers
powers = c(c(1:10), seq(from = 12, to=20, by=2))
# Call the network topology analysis function
sft = pickSoftThreshold(datExpr, powerVector = powers, verbose = 5)
# Plot the results:
sizeGrWindow(9, 5)
par(mfrow = c(1,2));
cex1 = 0.9;
# Scale-free topology fit index as a function of the soft-thresholding power
plot(sft$fitIndices[,1], -sign(sft$fitIndices[,3])*sft$fitIndices[,2],
xlab="Soft Threshold (power)",ylab="Scale Free Topology Model Fit,signed R^2",type="n",
main = paste("Scale independence"))
text(sft$fitIndices[,1], -sign(sft$fitIndices[,3])*sft$fitIndices[,2],
labels=powers,cex=cex1,col="red")
# this line corresponds to using an R^2 cut-off of h
abline(h=0.90,col="red")
```


```{r mean connectivity, echo=FALSE}
# Mean connectivity as a function of the soft-thresholding power
plot(sft$fitIndices[,1], sft$fitIndices[,5],
xlab="Soft Threshold (power)",ylab="Mean Connectivity", type="n",
main = paste("Mean connectivity"))
text(sft$fitIndices[,1], sft$fitIndices[,5], labels=powers, cex=cex1,col="red")
```


```{r clustrering, echo=FALSE}

net = blockwiseModules(datExpr, power = 6,
TOMType = "unsigned", minModuleSize = 30,
reassignThreshold = 0, mergeCutHeight = 0.25,
numericLabels = TRUE, pamRespectsDendro = FALSE,
saveTOMs = TRUE,
maxBlockSize = 10000,
saveTOMFileBase = "Seq-physioTOM",
verbose = 3)


# open a graphics window
sizeGrWindow(12, 9)
# Convert labels to colors for plotting
mergedColors = labels2colors(net$colors)
# Plot the dendrogram and the module colors underneath
plotDendroAndColors(net$dendrograms[[1]], mergedColors[net$blockGenes[[1]]],
"Module colors",
dendroLabels = FALSE, hang = 0.03,
addGuide = TRUE, guideHang = 0.05)

moduleLabels = net$colors
moduleColors = labels2colors(net$colors)
MEs = net$MEs;
geneTree = net$dendrograms[[1]];
save(MEs, moduleLabels, moduleColors, geneTree,
file = "networkConstruction-auto.RData")

#w.out grey


```


```{r gene-physio correlation, echo=FALSE, message=FALSE}

# Display the current working directory
getwd();
# If necessary, change the path below to the directory where the data files are stored.
# "." means current directory. On Windows use a forward slash / instead of the usual \.
workingDir = ".";
setwd(workingDir);
# Load the WGCNA package
library(WGCNA)
# The following setting is important, do not omit.
options(stringsAsFactors = FALSE);
# Load the expression and trait data saved in the first part
lnames = load(file = "input.RData");
#The variable lnames contains the names of loaded variables.
lnames
# Load network data saved in the second part.
lnames = load(file = "networkConstruction-auto.RData");
lnames

# Define numbers of genes and samples
nGenes = ncol(datExpr);
nSamples = nrow(datExpr);
# Recalculate MEs with color labels
MEs0 = moduleEigengenes(datExpr, moduleColors)$eigengenes
MEs = orderMEs(MEs0)
moduleTraitCor = cor(MEs, datTraits, use = "p");
moduleTraitPvalue = corPvalueStudent(moduleTraitCor, nSamples);

sizeGrWindow(10,6)
# Will display correlations and their p-values
textMatrix = paste(signif(moduleTraitCor, 2), "\n(",
signif(moduleTraitPvalue, 1), ")", sep = "");
dim(textMatrix) = dim(moduleTraitCor)
par(mar = c(6, 8.5, 3, 3));
# Display the correlation values within a heatmap plot
labeledHeatmap(Matrix = moduleTraitCor,
xLabels = names(datTraits),
yLabels = names(MEs),
ySymbols = names(MEs),
colorLabels = FALSE,
colors = greenWhiteRed(50),
textMatrix = textMatrix,
setStdMargins = FALSE,
cex.text = 0.5,
zlim = c(-1,1),
main = paste("Module-trait relationships"))
```


```{r defining individual varaibles}

# Define variable weight containing the weight column of datTrait
weight = as.data.frame(datTraits$PTW);
names(weight) = "PTW"
# names (colors) of the modules
modNames = substring(names(MEs), 3)
geneModuleMembership = as.data.frame(cor(datExpr, MEs, use = "p"));
MMPvalue = as.data.frame(corPvalueStudent(as.matrix(geneModuleMembership), nSamples));
names(geneModuleMembership) = paste("MM", modNames, sep="");
names(MMPvalue) = paste("p.MM", modNames, sep="");
geneTraitSignificance = as.data.frame(cor(datExpr, weight, use = "p"));
GSPvalue = as.data.frame(corPvalueStudent(as.matrix(geneTraitSignificance), nSamples));
names(geneTraitSignificance) = paste("GS.", names(weight), sep="");
names(GSPvalue) = paste("p.GS.", names(weight), sep="");

module = "green"
column = match(module, modNames);
moduleGenes = moduleColors==module;
sizeGrWindow(7, 7);
par(mfrow = c(1,1));
verboseScatterplot(abs(geneModuleMembership[moduleGenes, column]),

abs(geneTraitSignificance[moduleGenes, 1]),
xlab = paste("Module Membership in", module, "module"),
ylab = "Gene significance for PTW",
main = paste("Module membership vs. gene significance\n"),
cex.main = 1.2, cex.lab = 1.2, cex.axis = 1.2, col = module)


names(datExpr)

names(datExpr)[moduleColors=="green"]


geneInfo0 = data.frame(
geneTraitSignificance,
GSPvalue)
# Order modules by their significance for weight
modOrder = order(-abs(cor(MEs, weight, use = "p")));
# Add module membership information in the chosen order
for (mod in 1:ncol(geneModuleMembership))
{
oldNames = names(geneInfo0)
geneInfo0 = data.frame(geneInfo0, geneModuleMembership[, modOrder[mod]],
MMPvalue[, modOrder[mod]]);
names(geneInfo0) = c(oldNames, paste("MM.", modNames[modOrder[mod]], sep=""),
paste("p.MM.", modNames[modOrder[mod]], sep=""))
}
# Order the genes in the geneInfo variable first by module color, then by geneTraitSignificance
geneInfo0 = as.data.frame(geneInfo0)
geneOrder = order(geneInfo0$p.MM.green, -abs(geneInfo0$p.GS.PTW));
geneInfo = geneInfo0[geneOrder, ]

write.csv(geneInfo, file = "geneInfo.csv")


```


```{r blank}

```


#GO analysis, doesnt work

#```{r gene-ontology}
# Display the current working directory
getwd();
# If necessary, change the path below to the directory where the data files are stored.
# "." means current directory. On Windows use a forward slash / instead of the usual \.
workingDir = ".";
setwd(workingDir);
# Load the WGCNA package
library(WGCNA)
# The following setting is important, do not omit.
options(stringsAsFactors = FALSE);
# Load the expression and trait data saved in the first part
lnames = load(file = "input.RData");
#The variable lnames contains the names of loaded variables.
lnames
# Load network data saved in the second part.
lnames = load(file = "networkConstruction-auto.RData");
lnames

intModules = c("brown", "red", "salmon")
allLLIDs = datExpr
for (module in intModules)
{
# Select module probes
modGenes = (moduleColors==module)
# Get their entrez ID codes
modLLIDs = allLLIDs[modGenes];
# Write them into a file
fileName = paste("LocusLinkIDs-", module, ".txt", sep="");
write.table(as.data.frame(modLLIDs), file = fileName,
row.names = FALSE, col.names = FALSE)
}
# As background in the enrichment analysis, we will use all probes in the analysis.
fileName = paste("LocusLinkIDs-all.txt", sep="");
write.table(as.data.frame(allLLIDs), file = fileName,
row.names = FALSE, col.names = FALSE)


GOenr = GOenrichmentAnalysis(moduleColors, allLLIDs, organism = "chicken", nBestP = 10);

tab = GOenr$bestPTerms[[4]]$enrichment
write.table(tab, file = "GOEnrichmentTable.csv", sep = ",", quote = TRUE, row.names = FALSE)


```{r heatmap-prep}
# Display the current working directory
getwd();
# If necessary, change the path below to the directory where the data files are stored.
# "." means current directory. On Windows use a forward slash / instead of the usual \.
workingDir = ".";
setwd(workingDir);
# Load the WGCNA package
library(WGCNA)
# The following setting is important, do not omit.
options(stringsAsFactors = FALSE);
# Load the expression and trait data saved in the first part
lnames = load(file = "input.RData");
#The variable lnames contains the names of loaded variables.
lnames
# Load network data saved in the second part.
lnames = load(file = "networkConstruction-auto.RData");
lnames
nGenes = ncol(datExpr)
nSamples = nrow(datExpr)

```


```{r visualising_all}
# Calculate topological overlap anew: this could be done more efficiently by saving the TOM
# calculated during module detection, but let us do it again here.
dissTOM = 1-TOMsimilarityFromExpr(datExpr, power = 6, verbose = 3);
# Transform dissTOM with a power to make moderately strong connections more visible in the heatmap
plotTOM = dissTOM^7;
# Set diagonal to NA for a nicer plot
diag(plotTOM) = NA;
# Call the plot function
sizeGrWindow(9,9)
TOMplot(plotTOM, geneTree, moduleColors, main = "Network heatmap plot, all genes")
```


```{r visualising_top100}
nSelect = 100
# For reproducibility, we set the random seed
set.seed(10);
select = sample(nGenes, size = nSelect);
selectTOM = dissTOM[select, select];
# There’s no simple way of restricting a clustering tree to a subset of genes, so we must re-cluster.
selectTree = hclust(as.dist(selectTOM), method = "average")
selectColors = moduleColors[select];
# Open a graphical window
sizeGrWindow(9,9)
# Taking the dissimilarity to a power, say 10, makes the plot more informative by effectively changing
# the color palette; setting the diagonal to NA also improves the clarity of the plot
plotDiss = selectTOM^7;
diag(plotDiss) = NA;
TOMplot(plotDiss, selectTree, selectColors, main = "Network heatmap plot, selected genes")

```


```{r recalculating modules}
# Recalculate module eigengenes
MEs = moduleEigengenes(datExpr, moduleColors)$eigengenes
# Isolate weight from the clinical traits
weight = as.data.frame(datTraits$PTW);
names(weight) = "PTW"
# Add the weight to existing module eigengenes
MET = orderMEs(cbind(MEs, weight))
# Plot the relationships among the eigengenes and the trait
sizeGrWindow(5,7.5);
par(cex = 0.9)
plotEigengeneNetworks(MET, "", marDendro = c(0,4,1,2), marHeatmap = c(3,4,1,2), cex.lab = 0.8, xLabelsAngle
= 90)


```


```{r plotdendroggrtams}

# Plot the dendrogram
sizeGrWindow(6,6);
par(cex = 1.0)
plotEigengeneNetworks(MET, "Eigengene dendrogram", marDendro = c(0,4,2,0),
plotHeatmaps = FALSE)
# Plot the heatmap matrix (note: this plot will overwrite the dendrogram plot)
par(cex = 1.0)
plotEigengeneNetworks(MET, "Eigengene adjacency heatmap", marHeatmap = c(3,4,2,2),
plotDendrograms = FALSE, xLabelsAngle = 90)

```


```{r cytoscape}

# Recalculate topological overlap if needed
TOM = TOMsimilarityFromExpr(datExpr, power = 6);
# Read in the annotation file
# Select modules
modules = c("pink");
# Select module probes
probes = names(datExpr)
inModule = is.finite(match(moduleColors, modules));
modProbes = probes[inModule];
# Select the corresponding Topological Overlap
modTOM = TOM[inModule, inModule];
dimnames(modTOM) = list(modProbes, modProbes)
# Export the network into edge and node list files Cytoscape can read
cyt = exportNetworkToCytoscape(modTOM,
edgeFile = paste("CytoscapeInput-edges-", paste(modules, collapse="-"), ".txt", sep=""),
nodeFile = paste("CytoscapeInput-nodes-", paste(modules, collapse="-"), ".txt", sep=""),
weighted = TRUE,
threshold = 0.02,
nodeNames = modProbes,
nodeAttr = moduleColors[inModule]);
```
