## Supplementary Figures 1-8 for "*FSH*β links photoperiodic signalling to seasonal reproduction in Japanese quail"

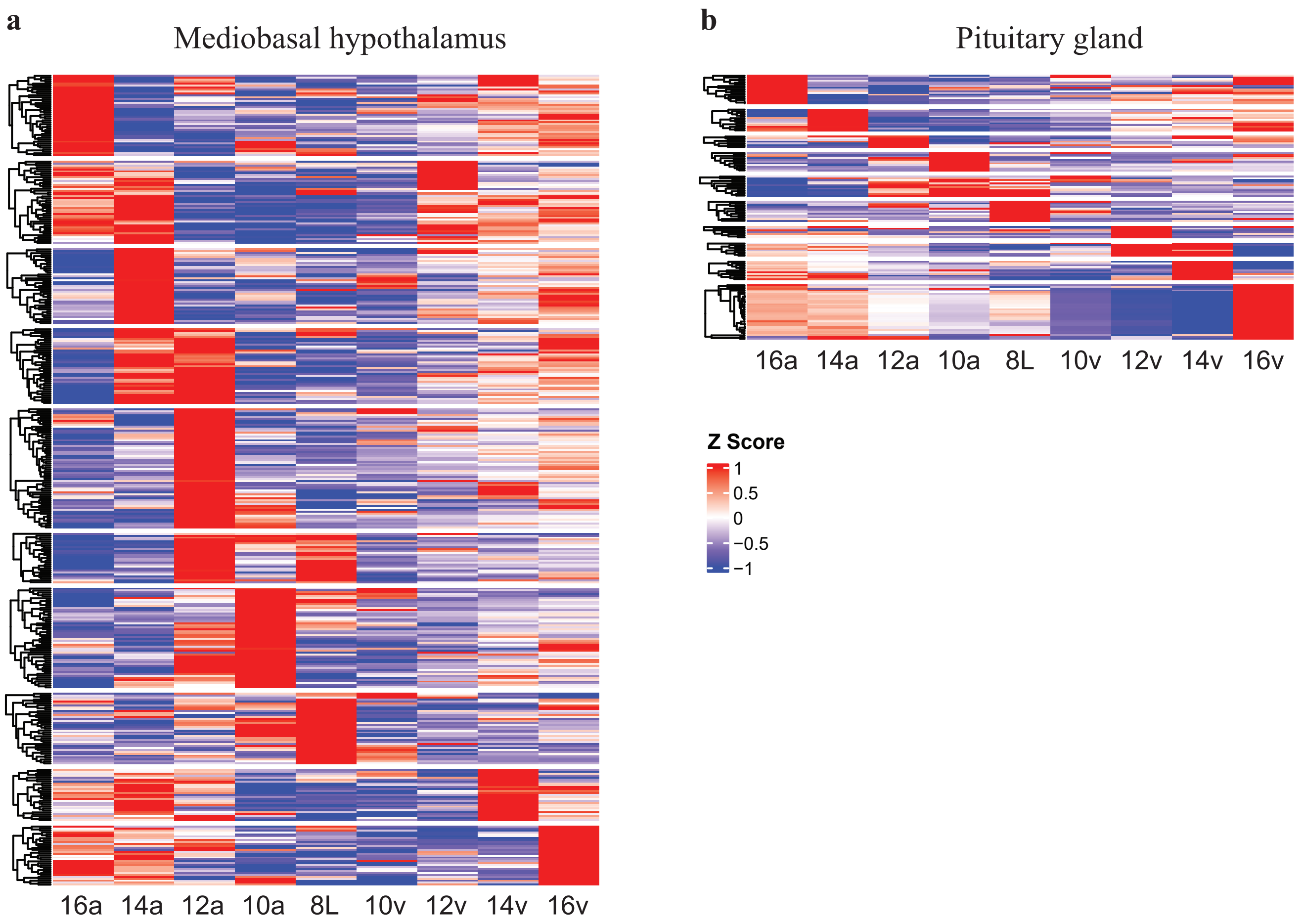


BioDare2.0 generated heat maps representing the **a** 398 transcripts found to be significant in the mediobasal hypothalamus and **b** 130 transcripts identified to be significant in the pituitary gland. Scale bar denotes significantly upregulated (red) and downregulated (blue) transcripts that display either rhythmic or spike patterns.

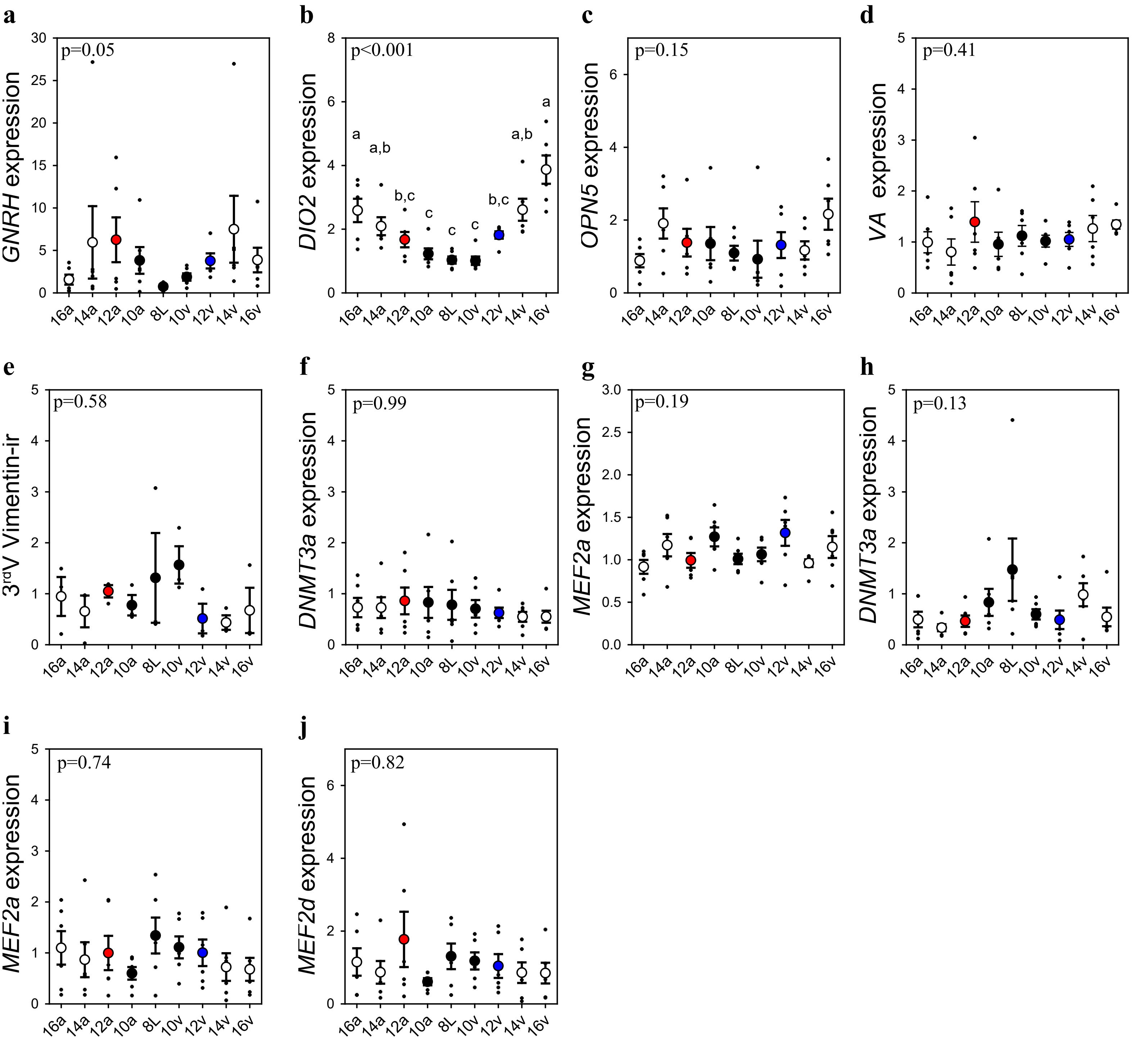


**a** *GNRH* expression in the preoptic area had a trend to significance suggesting low levels in 8L birds. **b-g** mediobasal hypothalamic expression of deiodinase type-2 (*DIO2*), photoreceptors (neuropsin [*OPN5*], vertebrate ancient opsin [*VA*]), tanycytes (vimentin) immunoreactivity along the third ventricle (3rdV), DNA methyltransferase 3a (*DNMT3a*) and transcription binding proteins myocyte enhancer factor 2a (*MEF2a*). **h-j,** Pituitary transcript expression for *DNMT3a* *MEF2a,* and *MEF2d*. One-way ANOVA were conducted to detected significance across the photoperiod treatment groups. Post-hoc Tukey test was conducted after Bonferroni correction to identify significant pairwise comparisons. Letters denote significant difference between photoperiod groups ($P<0.05$). Raw data available in Table S1.

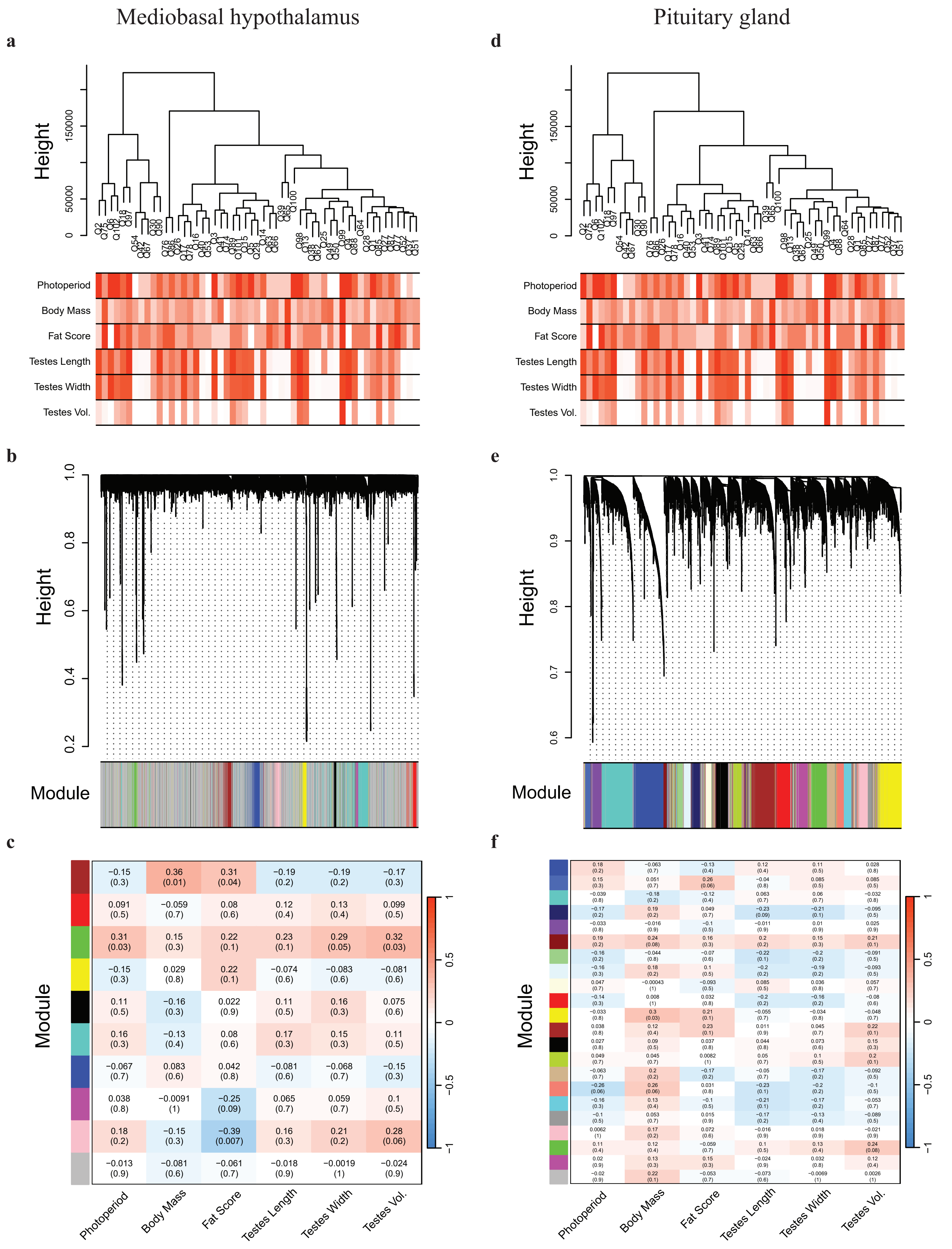


Weighted gene co-expression network analyses of the MBH and pituitary gland transcripts. The MBH **a** dendrogram and trait heatmap, **b** cluster dendrogram, and **c** module-trait relationships. The pituitary gland **d** dendrogram and trait heatmap, **e** cluster dendrogram, and **f** module-trait relationships.

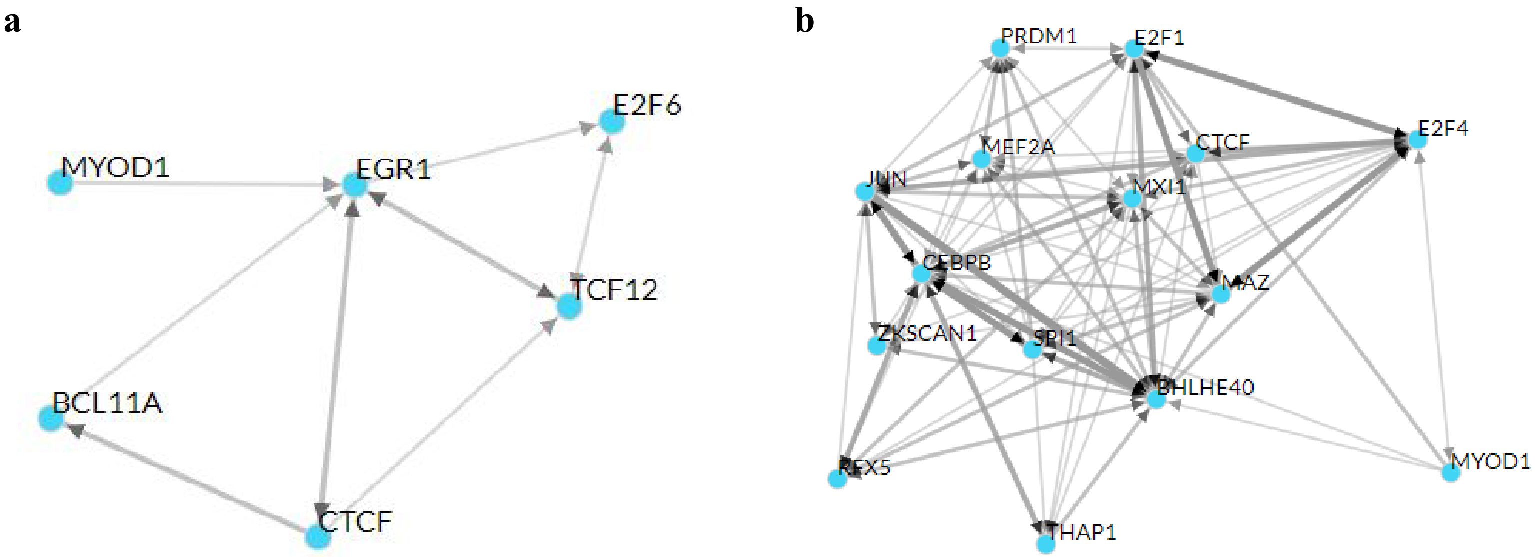


CHiP-X enrichment analyses to identify DNA binding motifs common across transcripts detected as significantly different based on BioDare2.0 analyses of the mediobasal hypothalamus (**a**) and pituitary gland (**b**). Thicker lines indicate stronger association across transcription factors. MBH had low commonality among transcripts and several showed no association. Pituitary gland transcripts had strong association among a relative few transcription factors. These findings indicate strong tissue-specific recruitment of transcription factors to drive photoperiod induced changes in transcript expression. Moreover, the strong association in the pituitary gland suggests few transcripts might orchestrate the seasonal change in expression. Abbreviations: BAF chromatin remodelling complex subunit 11A (BCL11A), Basic Helix-Loop-Helix family member E40 (BHLHE40), CCCTC-binding factor (CTCF), CCAAT Enhancer binding protein Beta (CEBPB), E2F Transcription Factor 1, (E2F1), E2F Transcription Factor 4 (E2F4), E2F Transcription Factor 6 (E2F6), Early growth response 1 (EGR1), Jun Proto-Oncogene (JUN), MAX Interactor 1 (MXI1), MYC Associated zinc finger protein (MAZ), Myocyte Enhancer  Factor 2A (MEF2A), Myoblast determination protein 1 (MYOD1), NK6 homeobox 2 (NKX62), PR domain zinc finger protein 1 (PRDM1), Regulatory factor X (RFX5), Spi-1 Proto-Oncogene (SPI1), Transcription factor 12 (TCF12), THAP Domain Containing 1 (THAP1), Zinc Finger With KRAB And SCAN Domains 1 (ZKSPAN1).

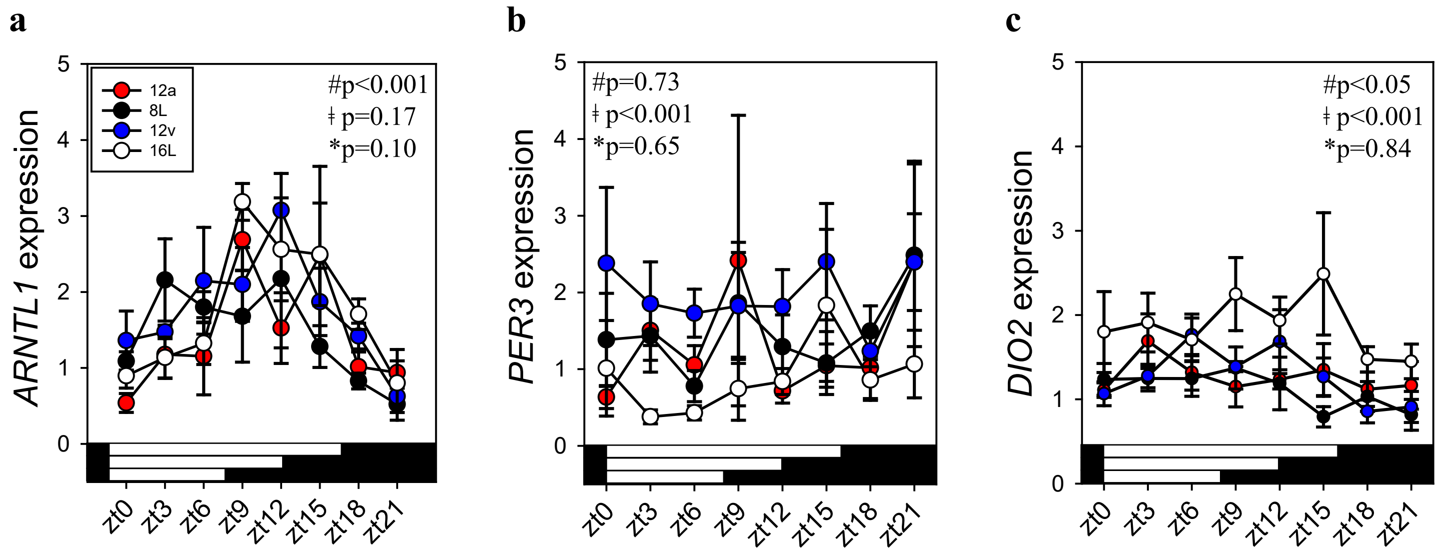


Circadian clock gene (*ARNTL1*, *PER3*) in the mediobasal hypothalamus from Japanese quail collected every 3 hours under short photoperiod (8L), autumnal equinox (12a; 12L:12D), vernal equinox (12v; 12L:12D) and long photoperiod (16L). **a** *ARNTL1* had a significant time of day effect, but no significant photoperiod effect or significant interaction. **b** There was a significant effect of photoperiod on *PER3* expression, but no significant time of day, or photoperiod treatment **c** thyroid hormone enzyme deiodinase type-2 (*DIO2*) expression in the quail MBH was observed to have a significant photoperiod effect and time of day, but no significant interaction. Two-way ANOVA was conducted for *ARNTL1*, *PER3,* and *DIO2* expression. ǂ indicates significant photoperiod treatment effect; # denotes significant time of day effect; * indicates significant interaction. Plot are mean +/- SEM and horizontal white and black bars indicate lights ON or OFF, respectively. Rhythmic analyses are presented in Extended Data Table 8 and raw data in Table S1.

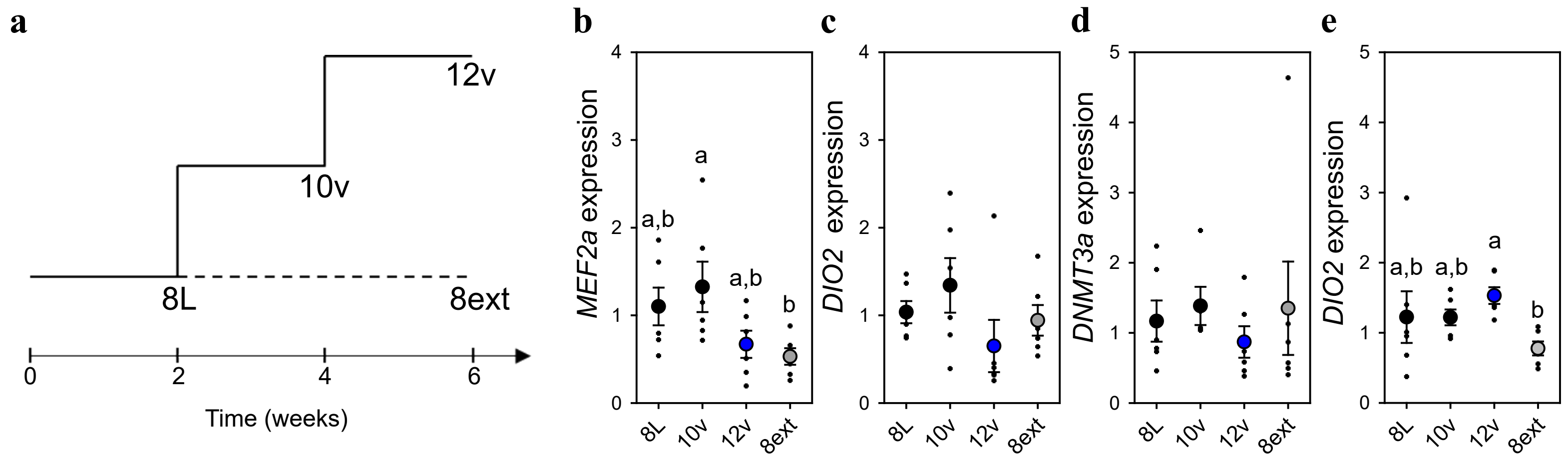


**a** Schematic representation of the experimental timline. A subset of quail were collected from short photoperiod (8L) and another group were kept on short photoperiod for an additional 4 weeks (8ext). Similar to the previous studies, another group of birds were transfered to 10L:14D for 2 weeks and another subset of birds were collected (10v). The final group of birds were transfered to 12L:12D for 2 weeks and served as the spring equinox group (12v). Quantitative PCR assays for *MEF2a*, *DIO2*, and *DNMT3a* expression in male quail housed in short photoperiod, spring transition to 10hr light (10v), the spring equinox (12v), or in short photoperiods for an extended 4 weeks (8ext). b One-way ANOVA revealed *MEF2a* expression in the pituitary gland varied significantly across the photoperiodic conditions ($F=3.37$, $P<0.05$) and Tukey's test identified higher levels in 10v compared to 8ext. **c** There was no significant difference in *DIO2* expression in the pituitary gland ($F=2.83$, $P=0.06$). **e** Pituitary gland *DNMT3a* did not vary significantly across photoperiodic conditions ($F=0.68$, $P=0.57$). *DIO2* in the medio-basal hypothalamus was significantly different across photoperiods ($F=3.19$, $P<0.05$). Tukey's test identified that 12v had higher levels compared to 8ext. Plot are mean +/- SEM with residual dot plots. Raw data are available in Table S1.

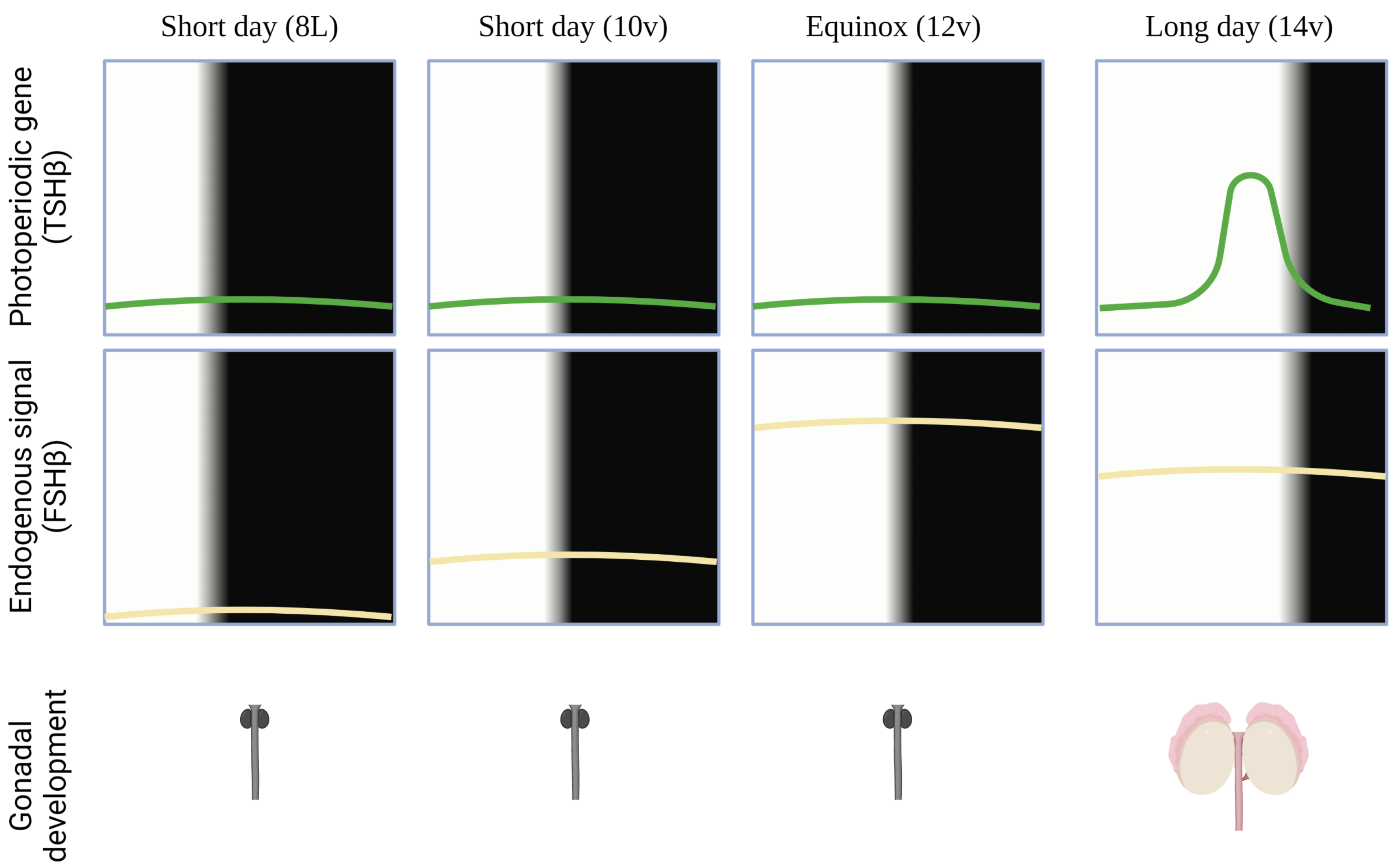


Two distinct pituitary cell types are involved in the external coincidence model for the avian photoperiodic response. Short photoperiods activate an endogenously generated programmed to increase *FSHβ* expression in the pars distalis of the pituitary gland. A gradual increase in non-stimulatory photoperiods, such as 10v and 12v, establish the photosensitive state and characterised by constituently expressed *FSHβ*. Photoperiods that extend beyond the critical day length (e.g., >12v) activates thyrotropes *TSHβ* expression in the pars tuberalis of the pituitary gland. The coincidence timing of long day *TSHβ* with the short day photosensitivity induced by increased *FSHβ* expression results in gonadal development. Figure was created using BioRender.

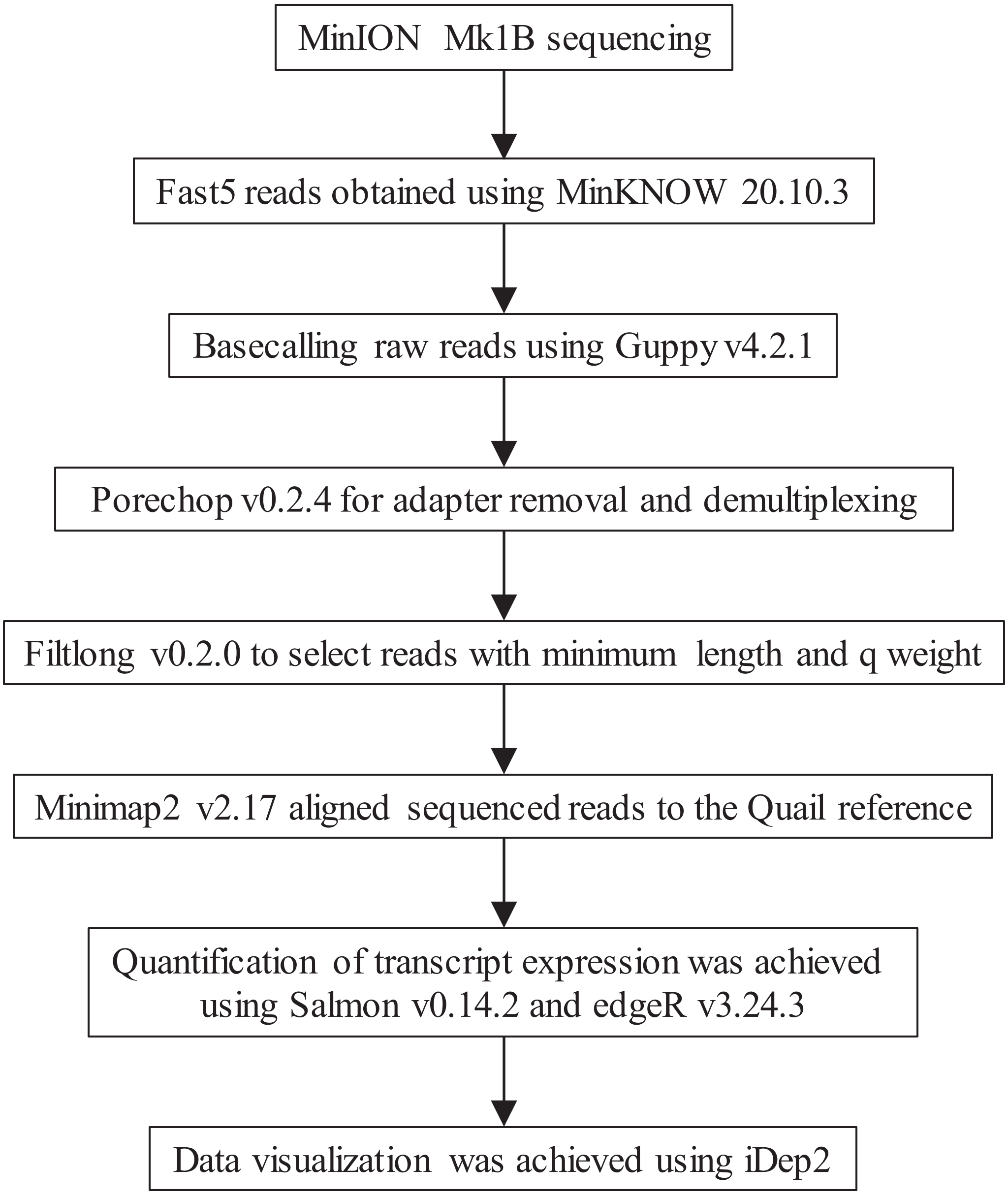
MinION Mk1B sequencing
Fast5 reads obtained using MinKNOW 20.10.3
Basecalling raw reads using Guppy v4.2.1
Porechop v0.2.4 for adapter removal and demultiplexing
Filtlong v0.2.0 to select reads with minimum length and q weight
Minimap2 v2.17 aligned sequenced reads to the Quail reference
Quantification of transcript expression was achieved using Salmon v0.14.2 and edgeR v3.24.3
Data visualization was achieved using iDep2
